# Opto-seq reveals input-specific immediate early gene induction in ventral tegmental area cell types

**DOI:** 10.1101/2023.06.22.546124

**Authors:** Rhiana C. Simon, Mary C. Loveless, Joshua X. Yee, Koichi Hashikawa, Garret D. Stuber, Larry S. Zweifel, Marta E. Soden

## Abstract

The ventral tegmental area (VTA) is a critical node in circuits governing motivated behavior and is home to diverse populations of neurons that release dopamine, GABA, glutamate, or combinations of these neurotransmitters. The VTA receives inputs from many brain regions, but a comprehensive understanding of input-specific activation of VTA neuronal subpopulations is lacking. To address this, we combined optogenetic stimulation of select VTA inputs with single-nucleus RNA sequencing (snRNAseq) and highly multiplexed *in situ* hybridization to identify distinct neuronal clusters and characterize their spatial distribution and activation patterns. Quantification of immediate early gene (IEG) expression revealed that different inputs activated select VTA subpopulations, which demonstrated cell-type specific IEG programs. Within dopaminergic subpopulations IEG induction levels correlated with differential expression of ion channel genes. This new transcriptomics-guided circuit analysis reveals the diversity of VTA activation driven by distinct inputs and provides a resource for future analysis of VTA cell types.

## Introduction

The VTA plays a critical role in regulating motivated behavior by sending diverse projections to forebrain regions (Morales and Margolis, 2017). Dopamine-producing neurons respond to a variety of reinforcing stimuli (Schultz et al., 1997, Wise, 2004, Bromberg-Martin et al., 2010), and are the major output of this region. Glutamate- and GABA-releasing VTA neurons also impact motivated behavior via connections within the VTA and with other mesolimbic regions (Dobi et al., 2010, Tan et al., 2012, van Zessen et al., 2012, Yoo et al., 2016). Beyond the broad neurotransmitter-based categorizations of neurons within the VTA, recent studies have demonstrated genetic, spatial and functional heterogeneity within these populations (Lammel et al., 2011, Morales and Margolis, 2017, Heymann et al., 2020, Poulin et al., 2020, Phillips et al., 2022). There is also tremendous anatomical and functional diversity of synaptic inputs onto VTA neurons (Lammel et al., 2012, Watabe-Uchida et al., 2012, Beier et al., 2015, Faget et al., 2016, Soden et al., 2020). VTA dopamine neurons can be activated by direct glutamatergic excitation, GABAergic disinhibition (via inhibition of local GABA neurons), and/or signaling by neuropeptides, which are co-released from many glutamatergic and GABAergic inputs (van den Pol, 2012).

Activation or inhibition of specific VTA inputs impacts distinct aspects of motivated behavior including action-outcome associations, reinforcement, and social preference (Lammel et al., 2012, Nieh et al., 2016, McHenry et al., 2017, Yang et al., 2018, Soden et al., 2020). Known VTA subpopulations also drive separable components of behavioral reinforcement (Heymann et al., 2020). The spatially restricted connectivity of specific VTA inputs, combined with the heterogeneity of behavioral regulation governed by both VTA inputs and outputs, suggests that distinct subpopulations within the VTA are recruited by different inputs to facilitate the many functions of the mesolimbic system. However, the genetic identity of these VTA cells and their responsivity to input-specific activation remains unresolved.

To address this question in an unbiased manner, we utilized an approach we have termed “opto-seq” that combines optogenetic stimulation of specific inputs followed by snRNA-seq of the mouse VTA, which allowed us to analyze the induction of IEGs and determine activation patterns within the VTA. We stimulated GABAergic inputs from the lateral hypothalamus (LH), which broadly innervate the VTA, GABAergic inputs from the nucleus accumbens (NAc), which strongly innervate the ventral VTA, and glutamatergic inputs from the medial prefrontal cortex (PFC), which also preferentially innervate the ventral VTA (Soden et al., 2020). Both LH and NAc GABAergic inputs synapse strongly onto VTA GABA neurons to disinhibit dopamine neurons, whereas PFC glutamatergic inputs synapse onto both dopamine and non-dopamine neurons, which likely results in direct activation and indirect feedforward excitation or inhibition (Bocklisch et al., 2013, Nieh et al., 2016, Soden et al., 2020).

Following optogenetic stimulation of these diverse inputs, we isolated and sequenced nuclei from over 40,000 VTA cells, including over 9,300 neurons. We mapped the spatial distribution of cell-type-specific molecular markers using highly multiplexed florescence *in situ* hybridization (FISH). Using both sequencing and FISH to assess IEG expression, we found that inputs selectively engaged distinct ensembles of cells and that patterns of stimulus-induced IEG expression were cell-type dependent. Among dopamine subpopulations, we identified sets of differentially expressed ion channel genes that correlated with IEG expression levels, suggesting that genes modulating intrinsic cellular properties shape neuronal responses to input-specific activation. Altogether, the use of opto-seq elevates our understanding of VTA heterogeneity and IEG complexity and represents a broadly useful approach for resolving the input-output relationships of a complex neuronal circuit.

## Results

### snRNA-Seq of mouse VTA identifies genetically defined subpopulations

To perform transcriptomic analysis of the mouse VTA following optogenetic stimulation of specific inputs, we injected *Slc32a1*-Cre (*Vgat*-Cre) mice bilaterally in the LH or NAc, and *Slc17a7*-Cre (*Vglut1*-Cre) mice bilaterally in the PFC with a Cre-dependent virus encoding channelrhodopsin (AAV1-FLEX-ChR2-YFP). Control mice were injected with AAV1-FLEX-YFP, and all mice were implanted with bilateral fiber optic cannulas above the VTA. Following recovery from surgery and extensive acclimation to the patchcord, mice received 15 min of 20 Hz blue light stimulation. Immediately after stimulation the VTA was microdissected and snap frozen. VTA tissue from 3-4 mice per group was pooled and nuclei were isolated and sequenced using the 10x Genomics Next GEM platform (**Figure 1A**). For comprehensive transcriptomic analysis of VTA cell types, 41,468 total nuclei from all four groups underwent doublet removal, algorithmic integration and batch correction (Butler et al., 2018, Haghverdi et al., 2018), followed by graph-based clustering via the Seurat R package (Stuart et al., 2019) (see Methods and **Figure S1A-G**). Clusters expressing neuronal markers were isolated and re-clustered, and nuclei that failed to meet quality control standards were removed (see Methods and **Figure S1H-K**). This resulted in a final population of 9,336 high-quality sequenced neurons used for subsequent analysis.

**Figure 1.**
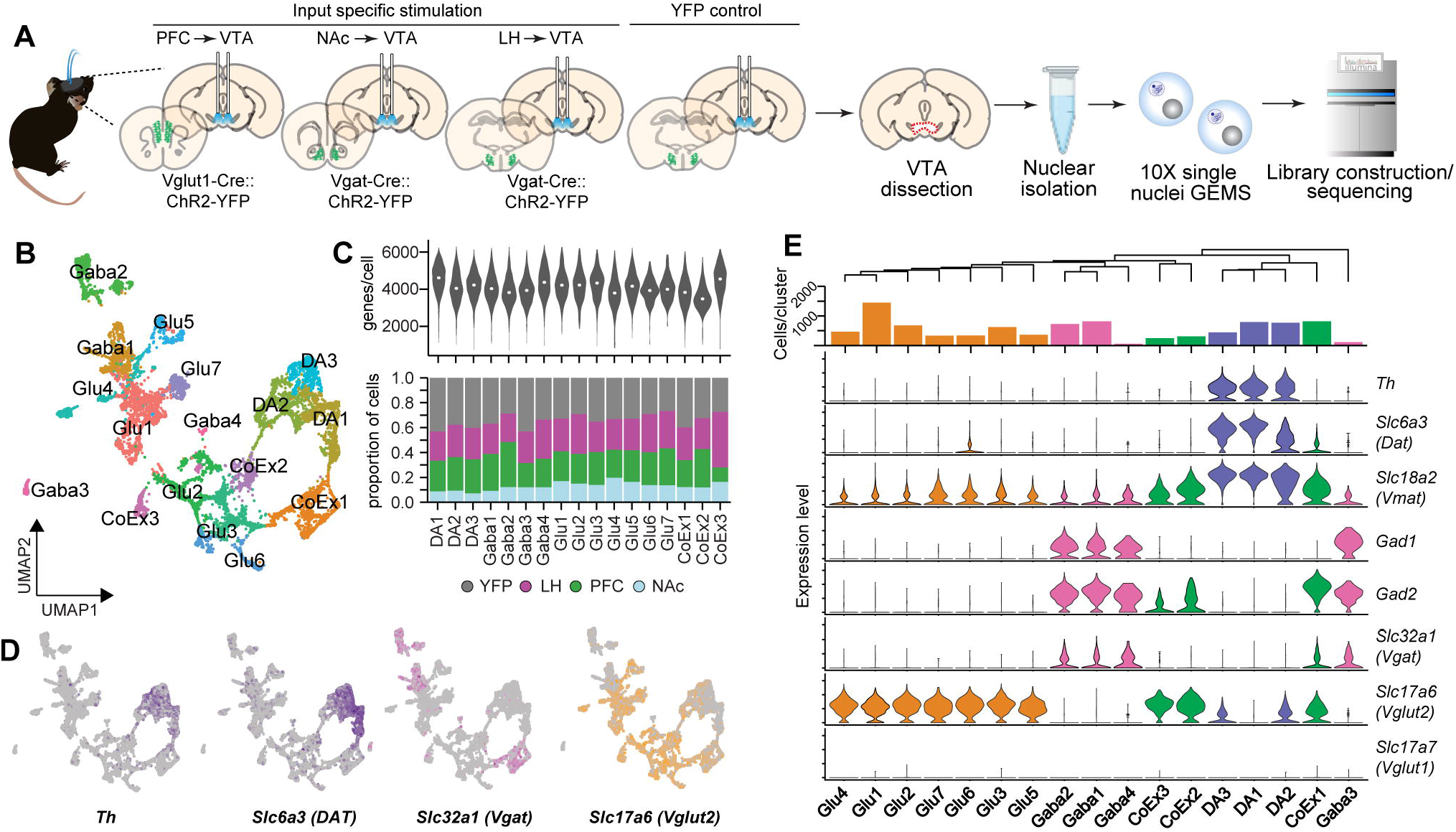
Opto-seq and identification of VTA neuronal subclusters. **A)** For input-specific stimulation, AAV1-FLEX-ChR2-YFP was injected in the PFC of Vglut1-Cre mice or the NAc or LH of Vgat-Cre mice and fiber optic cannulae were implanted above the VTA. Control mice received AAV1-FLEX-YFP. Following stimulation, the VTA was dissected and nuclei were isolated and sequenced. **B)** UMAP plot of neuronal subclusters. N=9,336 nuclei. **C)** Top: number of genes per cell across subclusters. Bottom: proportion of cells in each subcluster from each stimulus group. **D)** Feature plots showing expression of dopamine (*Th* and *Slc6a3*), GABA (*Slc32a1*), and glutamate (*Slc17a6*) cell type markers across subclusters. **E)** Top: dendrogram showing relationship between subclusters and bar graph showing number of cells/subcluster. Bottom: violin plots showing expression of cell type markers across subclusters.

VTA neurons segregated into 17 subclusters (**Figure 1B**). The number of genes per cell was consistent across clusters, and neurons from each stimulation condition were distributed evenly, though the NAc group had fewer cells overall than the other conditions (**Figure 1C**). The expression of neurotransmitter-related genes was used to manually label each cell type (**Figure 1D-E**). We identified 3 primarily dopaminergic clusters, dubbed DA1, DA2, and DA3, which expressed markers *Th, Slc6a3* (*Dat*), and *Slc18a2* (*Vmat*). DA2 and DA3 also showed low-level co-expression of the glutamatergic marker *Slc17a6* (*Vglut2*). We identified two large and two small GABA clusters (GABA1-GABA4) defined by expression of *Gad1, Gad2*, and *Slc32a1* (*Vgat*). Seven clusters were primarily glutamatergic (Glu1-7), as evidenced by high expression of *Vglut2*. Finally, we identified 3 clusters with co-expression of GABAergic and glutamatergic markers (CoEx1-3). All three showed expression of *Gad2*, but not *Gad1*, along with *Vglut2*, similar to a *Gad2*+/*Gad1*-population previously identified in the rat VTA (Phillips et al., 2022).

To determine if our tissue dissections contained cells from neighboring regions, we used the Allen Brain Atlas (Lein et al., 2007) to identify genes specifically expressed in regions that border the VTA. Most of these genes showed very low expression in our dataset, indicating relatively clean VTA dissections (**Figure S1L-O**). However, we did detect modest expression of some genes expressed in the supramammilary nucleus (ventral to the rostral VTA) in some Glu and CoEx clusters (**Figure S1L**). Overall, we were confident that our data represented VTA transcriptional heterogeneity, which allowed us to next evaluate input-specific modulation of VTA subtypes.

### Optogenetic stimulation drives cell type- and input-specific patterns of IEG activation

Neuronal activation triggers gene expression programs including rapid induction of IEGs such as *Fos, Egr1*, and *Jun*, which are then followed by secondary response genes that support long-term structural changes in neurons (Lanahan and Worley, 1998, Fowler et al., 2011, Okuno, 2011, Tyssowski et al., 2018). To evaluate stimulation-induced IEG patterns within the VTA we utilized a literature-derived inventory of 139 IEGs (Wu et al., 2017), of which 137 were detectable in our dataset. Given that most neuronal activation studies have utilized only a small subset of this IEG list, we also generated a shortlist of IEGs to test (*Arc, Bdnf, Crem, Egr1, Fos, Fosb, Fosl1, Fosl2, Homer1, Jun, Junb, Jund*, and *Myc*) that have been commonly used as markers of neuronal activity (Okuno, 2011, Tyssowski et al., 2018, Yap and Greenberg, 2018).

First, using the full IEG list and looking at the total population of neurons, we found that all stimulation conditions significantly increased expression of many IEGs compared to YFP control (PFC: 44 genes, NAc: 63, LH: 40), with few genes (1-5 per group) showing significantly reduced expression (**Figure 2A**). We next evaluated whether input driven IEG programs were variable between cell types or between inputs. To analyze IEG variability we collapsed the subclusters into four major cell classes (DA, Glu, GABA, and CoEx) separated by stimulus, and for each class calculated the change in expression (log2FC compared to YFP) for each IEG on the full list. A correlation analysis found that each cell class tended to cluster together regardless of input stimulation (**Figure 2B**), indicating that the pattern of IEG induction was more strongly determined by cell identity than by stimulation type. Indeed, the average correlation coefficient (Pearson’s *r*) for comparisons of same cell class between different inputs was significantly higher than for comparisons of the same input between different cell classes, or comparisons between groups not matched by class or input (**Figure 2C**). This relationship held true if correlation coefficients were calculated based only on the shortlist of commonly studied IEGs (**Figure 2C**).

**Figure 2.**
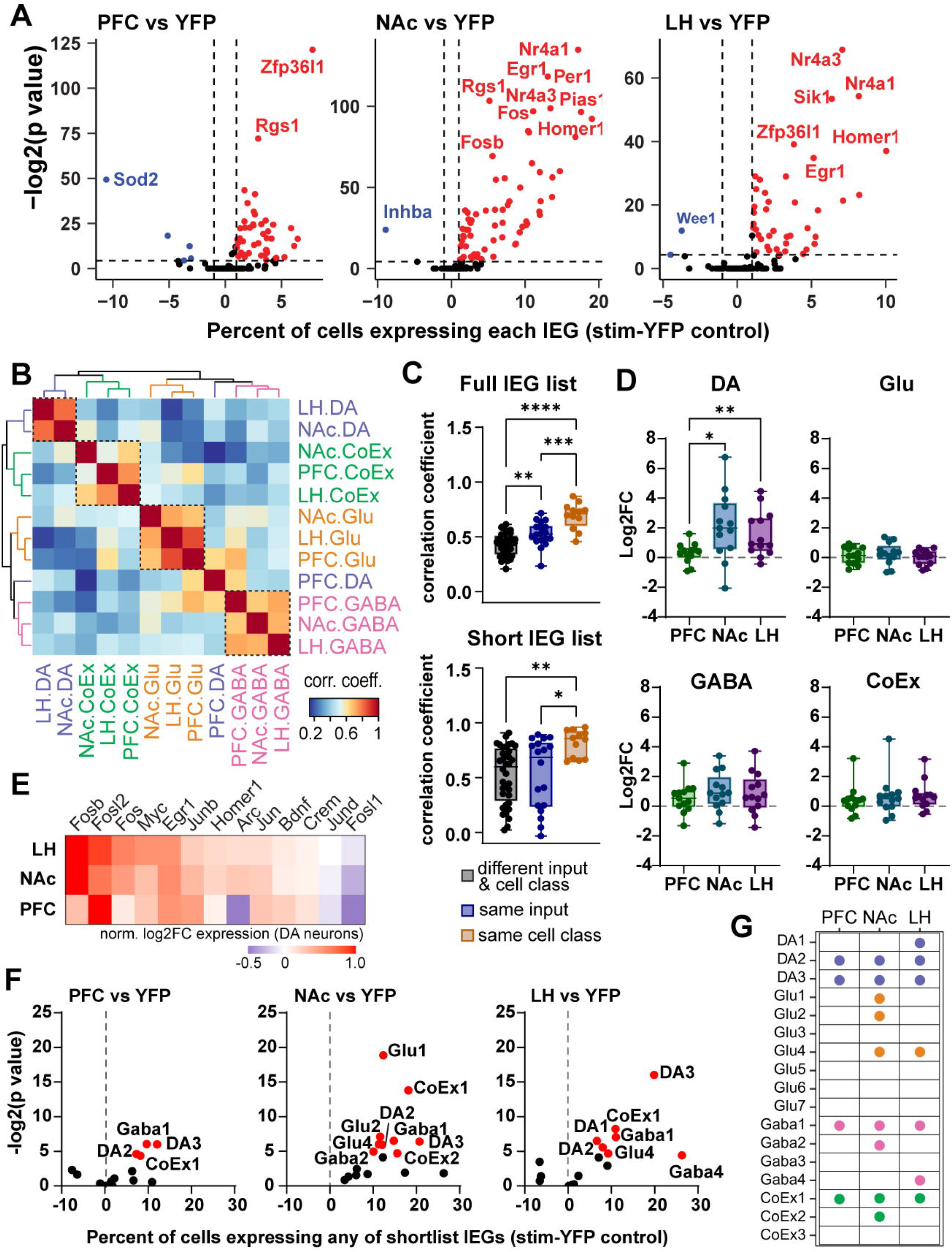
Optogenetic stimulation of distinct VTA inputs generates variable IEG expression. **A)** Volcano plots showing the percent of all neurons in each stimulation condition expressing each IEG minus YFP control, plotted against the -log2(P value) calculated by Fisher’s Exact Test. Genes with significantly increased expression (P<0.05) are in red, genes with significantly decreased expression are in blue. **B)** Heatmap showing the correlation coefficient (Pearson’s *r*) of the log2FC in expression of each IEG (full list) compared to YFP control, comparing between cell types and stimuli. **C)** Correlation coefficients for comparisons between pairs of different input and cell type, the same input but different cell type, or the same cell type but different input. Top: full IEG list, One-way ANOVA *F*_*(2*,*63)*_=27.50, P<0.0001. Bottom: short IEG list, One-way ANOVA *F*_*(2*,*63)*_*=5*.*922*, P=0.0044. **D)** Log2FC compared to YFP of shortlist IEGs, separated by cell class and input. DA: One-way RM ANOVA *F*_*(1*.*200*, *14*.*40)*_ = 11.15, P=0.0034. **C&D**: Box and whisker plots depict median, 25^th^/75^th^ percentiles, and min/max. Tukey’s multiple comparisons *P<0.05,**P<0.01,***P<0.001, ****P<0.0001. **E)** Log2FC in expression of each shortlist IEG in DA neurons, normalized to the maximum for each input. **F)** For each input, the percent of cells within each subcluster expressing any of shortlist IEGs (minus YFP control), plotted against the-log2(P value) calculated by Fisher’s Exact Test. Clusters with significantly increased expression (P<0.05) are in red. **G)** Summary plot of significantly activated clusters from **F**.

One notable exception to the rule of cell classes clustering together was DA neurons, as the IEG expression pattern induced by PFC inputs did not cluster with the expression induced by NAc or LH inputs (**Figure 2B**). To further examine this, we plotted the log2FC in expression (compared to YFP) of each shortlist IEG for each cell class, separated by input. For Glu, GABA, and CoEx classes there was no significant difference in the average change in expression of this gene set between inputs, but for DA neurons both LH and NAc inputs generated significantly higher IEG induction than did PFC inputs (**Figure 2D**). However, even when FC values for IEG expression in DA neurons were normalized within each input we observed that induction patterns were highly similar between NAc and LH stimulation, but were less similar between these groups and PFC stimulation (**Figure 2E**). Together, these data indicate that cell type is an important determinant of IEG expression patterns, but that PFC to VTA glutamatergic stimulation induces both weaker IEG induction and different IEG programs in dopamine neurons compared to LH- and NAc GABAergic disinhibition.

Finally, to identify specific neuronal clusters activated by each input, we again focused on the shortlist of highly studied IEGs. For each stimulation group we plotted the change compared to YFP in the percent of cells in each cluster expressing at least 1 copy of any of these IEGs (**Figure 2F**) and found that each input significantly activated a unique set of clusters. Dopamine clusters DA2 and DA3 were activated by all 3 inputs, while DA1 was significantly activated only by LH inputs. LH inputs activated 1 glutamate cluster while NAc inputs activated 3 glutamate clusters, and all 3 inputs activated at least one GABA and one CoEx cluster (**Figure 2G**).

### Spatial resolution of VTA subpopulations with highly-multiplexed *in situ* hybridization

To select potential genetic markers that define neuronal subclusters, we identified genes that distinguish clusters from one another via conserved marker analysis (**Figure 3A**). To develop a comprehensive spatial profile for VTA cell types, we then selected 11 candidate cell-type markers based on their specificity in snRNAseq data and probe performance during *in situ* pilots. We also included 4 neurotransmitter markers (*Th, Gad1, Gad2*, and *Vglut2*; **Figure 3B**). Using sequential rounds of FISH (HCR, Molecular Instruments, followed by RNAScope, ACDBio), we probed for a panel of 15 total genes in 9 coronal VTA sections spanning from -2.92 mm to -3.88 mm from bregma (**Figure 3C-D**). As expected, *Th* was abundantly expressed, peaking in the central sections, and distributed broadly across the medial-lateral axis (**Figure 3E-H**). *Gad1* and *Gad2* were most abundant in the caudal VTA, but we observed a *Gad2+*/*Gad1-*population in the rostral-medial VTA. *Vglut2* neurons were most abundant in the rostral VTA, particularly medial and ventral to the *Th*+ populations. 79.7% of neurons expressed a single neurotransmitter phenotype (i.e. *Th* only, *Vglut2* only, or *Gad* only), but we also identified populations of co-expressing cells (**Figure 3F**). The majority (73.8%) of GABA cells that co-expressed *Vglut2* and/or *Th* were *Gad2* only cells, lacking *Gad1* expression.

**Figure 3.**
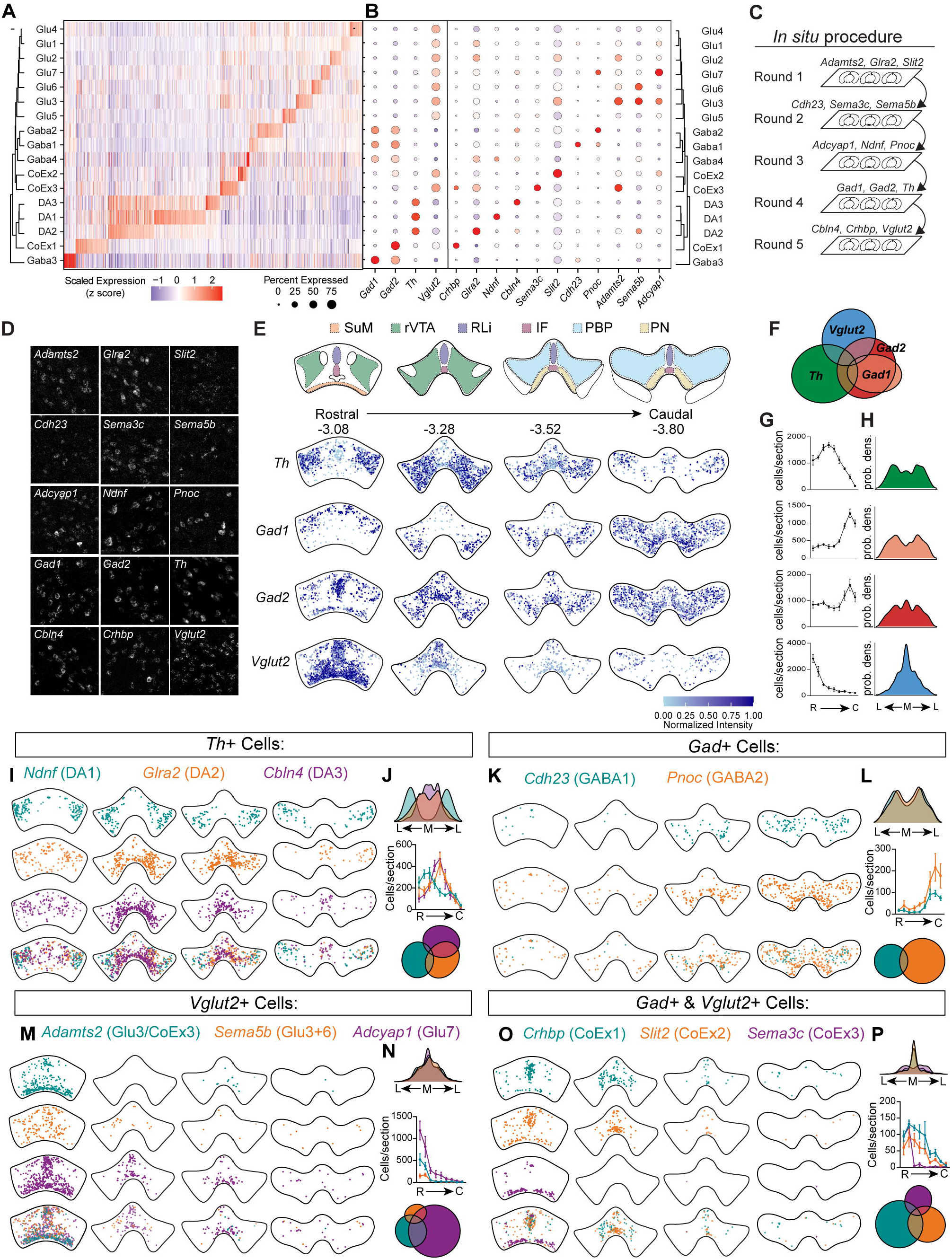
Spatial resolution of VTA subclusters. **A)** Heatmap showing scaled expression (Z scored) of differentially expressed genes across neuronal subclusters. **B)** Dot plot of genes selected for FISH, showing the percent of cells in each subcluster that express each gene (size of dot) and the z-scored expression (color). **C)** Schematic of 5 rounds of FISH performed on the same tissue sections. Rounds 1-4 were HCR, round 5 was RNAscope. **D)** Example FISH images for each gene. **E)** Top: diagram showing VTA subdivisions (SuM: supermammilary, rVTA: rostral VTA, RLi: rostrolinear nucleus, IF: interfascicular nucleus, PBP: parabrachial pigmented nucleus, PN: paranigral nucleus). Bottom, representative expression plots for 4 of 9 sections from one mouse. Each point represents one cell positive for the indicated gene; color reflects the normalized signal intensity. **F)** Euler plot combining data from 4 mice showing overlap in expression of canonical cell type markers. **G)** Average number of cells per section for each indicated gene, rostral (R) to caudal (C) (N=4 mice, 9 sections from bregma -2.92 mm to -3.88 mm, mean±SEM). **H)** Probability density plot showing medial (M) to lateral (L) distribution of cells. Data pooled from 4 mice, 9 sections per mouse. **I-P)** Quantification of cells positive for the indicated marker (*Th, Gad1* or *2, Vglut2*, or *Gad1* or *2* + *Vglut2*) as well as the indicated marker gene. **I, K, M and O)** Representative expression plots of positive cells in 4 of 9 sections from one mouse. **J, L, N and P)** Top: probability density plots showing medial-lateral distribution of positive cells, pooled across 4 mice. Middle: average cells/section from rostral to caudal, (N=4 mice, mean±SEM). Bottom, Euler plot combining data from 4 mice showing overlap between indicated genes.

Next, we examined the spatial distribution of our dopamine cluster markers, isolating only *Th*+ cells to increase specificity (**Figure 3I-J and Figure S2A-C**). DA1 marker *Ndnf* was detected in the lateral VTA (and was also present in the SNc, data not shown) and was most abundant in rostral sections. DA2 marker *Glra2* and DA3 marker *Cbln4* were found primarily in the central sections of the VTA and were more medial than *Ndnf*. Although we observed significant spatial overlap between these probes, we found that *Cbln4*+ cells were concentrated significantly more ventrally and medially than *Glra2*+ cells, quantified by the cumulative distributions of cells across the dorso-ventral and medial-lateral axes (**Figure S2D-E**). To better segregate the spatial organization of the DA2 and DA3 clusters, we probed additional sections from the same animals with two alternate candidate markers, *Lepr* (DA2) and *Grp* (DA3; **Figure S2F**). We saw minimal overlap between *Th*+ *Lepr* and *Grp* neurons and found that *Grp* was more restricted to the ventromedial VTA while *Lepr* was more dorsolateral, though not as lateral as *Ndnf* (**Figure S2G-J**). This organization of DA neurons into a rostral/lateral cluster (DA1), a ventromedial cluster (DA3), and an intermediate dorsolateral cluster (DA2) is consistent with the segregation of previously identified DA subtype markers across our DA clusters (Heymann et al., 2020, Poulin et al., 2020) (**Figure S2K**).

We next isolated cells that expressed *Cdh23* (GABA1) or *Pnoc* (GABA2) plus *Gad1* and/or *Gad2* (**Figure 3K-L and Figure S3A-B**). *Cdh23* and *Pnoc* GABA neurons were both enriched in the caudal VTA, but few cells co-expressed both markers (**Figure 3L**). In the most caudal sections, we observed a strong ventral band of *Pnoc*-expressing cells, while *Cdh23* cells were concentrated more dorsally. Our DA3 marker, *Cbln4*, also showed some expression in the GABA2 cluster (**Figure S3C**); when we isolated *Gad*+ *Cbln4*+ cells, they were located to the same caudal ventral band as *Pnoc* (**Figure S3D**), supporting the observed spatial separation of these GABA clusters.

Selecting specific marker genes for VTA glutamate neurons was challenging as most potential markers were expressed promiscuously across glutamate clusters. We selected *Adamts2, Sema5b*, and *Adcyap1* based on their relatively restricted expression and detectability via *in situ*. All three genes were primarily expressed in the rostral VTA, with *Adamts2* showing concentrated expression in the ventral supramammilary region, *Sema5b* showing distributed expression, and *Adcyap1* showing concentrated expression ventrally and along the midline (**Figure 3M-N and Figure S3E-G**).

Finally, to examine markers for CoEx clusters, we isolated cells that expressed *Vglut2* and either *Gad1* or *Gad2* (**Figure 3O-P and Figure S3H-J**). Among these cells, CoEx1 marker *Crhbp* was primarily expressed in the rostral VTA, concentrated on the midline. CoEx2 marker *Slit2* showed a similar distribution, with a concentration of cells in the interfascicular nucleus. CoEx3 marker *Sema3c* was specifically located to the supramammilary nucleus. Altogether, our *in situ* results provide a high-resolution account of the spatial distribution of genetically dissociable VTA cell types.

### Multiplex FISH analysis reveals variable IEG induction by innervation pattern, cell type, and input

To examine spatial expression of IEGs, particularly among dopamine neuron subtypes, we implanted and stimulated additional mice as described above and performed 3 rounds of FISH on 4 sections from each of 4 mice per group to label neurotransmitter markers *Th, Gad2*, and *Vglut2*, dopamine subgroup markers *Ndnf, Glra2*, and *Grp*, and IEGs *Egr1, Fos*, and *Homer1* (**Figure 4A**). These IEGs were selected based on their relatively high variability between conditions in our sequencing data (**Figure S4A**). We used alternative DA3 marker *Grp* as opposed to *Cbln4* to increase specificity. We also imaged ChR2-YFP fibers in the same sections. Like our previous studies we found that LH-GABA fibers broadly innervated the entire VTA with low variability and little dorsoventral or medial-lateral bias in fiber intensity **(Figure 4B-C and Figure S4B-C**). NAc-GABA inputs showed the most variability in fiber intensity, preferentially innervating the ventral and medial VTA (**Figure 4B-C and Figure S4B-C**). PFC-glutamate fibers also showed a ventral bias, though were not as variable as NAc inputs (**Figure 4B-C and Figure S4B-C**).

**Figure 4.**
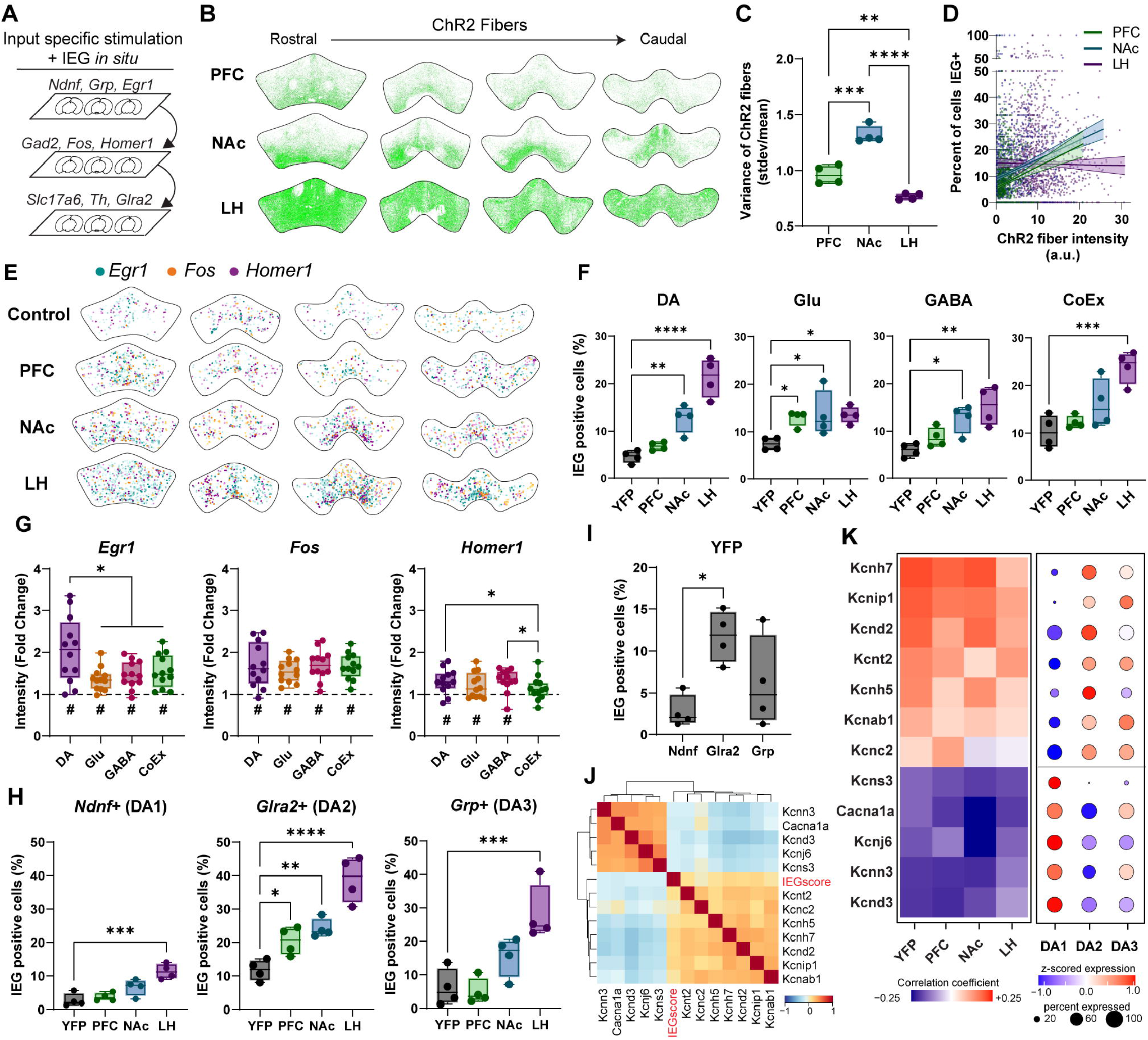
In situ analysis of IEG expression and identification of ion channel genes that correlate with IEG activity. **A)** Schematic of 3 rounds of FISH. Rounds 1 and 2 were HCR, round 3 was RNAscope. ChR2-YFP fibers were imaged during Round 1. **B)** Representative images of ChR2-YFP fibers from different inputs. **C)** Variance of fiber innervation, quantified as the standard deviation of fluorescence intensity across all cells divided by the mean (N=4 mice/group. One-way ANOVA *F*_*(2*,*9)*_*=66*.*34*, P<0.0001, Tukey’s multiple comparisons.). **D)** All sections were divided into a grid of approximately 150 μm × 150 μm squares. Plotted is a linear regression of average ChR2 fiber intensity (artificial units) within each square against the percent of cells positive for any of the 3 IEGs (Test of non-zero slope: PFC: *F*_*(1*,*1315)*_=35.40, P<0.0001, NAc: *F*_*(1*,*1436)*_=75.83, P<0.0001, LH *F*_*(1*,*1407)*_=0.1583, P=0.6907). **E)** Representative expression plots of cells positive for *Egr1, Fos*, or *Homer1* following stimulation of indicated inputs. **F)** Percent of each cell type positive for one or more IEG. (N=4 mice. One-way ANOVA, DA: *F*_*(3*,*12)*_=31.15, P=0.0001; Glu: *F*_*(3*,*12)*_=4.529, P=0.0241; GABA: *F*_*(3*,*12)*_=8.562, P=0.0026; CoEx *F*_*(3*,*12)*_=11.18, P=0.0009; Dunnett’s multiple comparisons, stim groups vs. YFP.) **G)** Fold change in fluorescence intensity of each IEG in positive cells relative to YFP control, pooled across all stimulus groups. (N=12 mice. One sample t-test of mean different than 1: #p<0.05. One-way RM ANOVA, *Egr1*: *F*_*(1*.*201*,*13*.*21)*_=10.79, P=0.0043; *Fos*: *F*_*(1*.*284*,*14*.*12)*_=0.7203, P=0.4443, *Homer1*: *F*_*(1*.*782*,*19*.*60)*_=5.359, P=0.0164. Tukey’s multiple comparisons. **H)** Percent of each cell type positive for one or more IEG. (N=4 mice/group. One-way ANOVA, *Ndnf*: F_(3,12)_=14.11, P=0.0003; *Glra2*: F_(3,12)_=25.32, P<0.0001, *Grp*: *F*_*(3*,*12)*_=11.93, P=0.0007; Dunnett’s multiple comparisons, stim groups vs. YFP.) **I)** Percent of Th+ cells of each type in the YFP group that express one or more IEGs (n=4 mice, one-way ANOVA F_(2,9)_= 5.652, P=0.0257, Tukey’s multiple comparisons ^*^P<0.05). All box and whisker plots depict median, 25^th^/75^th^ percentiles, and min/max. Forall post-hoc comparisons*P<0.05,^**^P<0.01,^***^P<0.001, ^****^P<0.0001. **J)** Heatmap showing the correlation coefficient (r) of select ion channel genes with the IEG score (summed IEG activity) in a given cell. Analysis was performed on sequencing data from all cells in clusters DA1, DA2, and DA3. **K)** Left: Heatmap of correlation coefficient of each ion channel gene with the IEG score, separated by stimulus condition. Right: dot plot showing percent of cells in each subcluster expressing each gene, and relative expression (z score) among DA clusters.

We overlayed a grid on each image and quantified fiber intensity and the percent of cells positive for any IEG within each division. IEG activation was significantly correlated with ChR2 fiber intensity of both NAc inputs and PFC inputs (**Figure 4D**). Consistent with this finding, for both NAc and PFC inputs a disproportionately high number of IEG positive cells were found in the divisions with the brightest ChR2 fibers (**Figure S4D-F**). The fiber intensity of LH inputs was uncorrelated with IEG activation, likely due to the overall uniformity of innervation (**Figure 4D and S4F**).

Using this FISH dataset we classified cells as DA, Glu, GABA, or CoEx based on expression of *Th, Gad2*, and *Vglut2* and determined the percent of cells of each type that were positive for any of the probed IEGs (**Figure 4E-F and S4G**). Stimulation of NAc and LH GABA inputs increased the percentage of IEG positive dopamine neurons and GABA neurons, while all three inputs increased IEG positive glutamate neurons. Only LH input stimulation significantly increased IEG positive CoEx neurons. We also examined expression of individual IEGs by quantifying the fold change in fluorescence intensity of each probe in the pooled stimulus groups compared to control (**Figure 4G**). *Egr1* and *Fos* were both significantly induced relative to control in all cell types, and *Homer1* was significantly induced in DA, Glu, and GABA, but not CoEx neurons. We observed a preferential activation of *Egr1* in dopamine neurons relative to other cell types; when separated by stimulus this pattern was largely consistent in NAc and LH inputs, but not in PFC inputs, which showed weaker induction (**Figure S4H**).

We then isolated *Th* positive neurons that expressed dopamine subgroup markers. Stimulation of LH-GABA inputs increased the percentage of IEG positive cells in all three subgroups, while NAc and PFC inputs increased the percentage of IEG positive *Glra2* neurons (DA2) only (**Figure 4H and S4G**). This was largely consistent with our sequencing experiment, apart from the DA3 subtype, where we had also seen activation by PFC and NAc inputs. This may reflect differences in sensitivity between the methods or in the number of IEGs examined, or may be because not all dopamine neurons are labeled by these marker genes (**Figure S4I**). Indeed, we found that NAc inputs significantly activated *Th* positive neurons that were unlabeled by any of our three markers (**Figure S4J**).

### Identification of ion channel genes correlated with IEG expression

In YFP control animals, a significantly higher percentage of *Glra2*+ cells were IEG positive at baseline compared to *Ndnf*+ cells (**Figure 4I**), with *Grp*+ cells showing more variable expression. *Glra2*+ cells also showed the highest rates of IEG induction in all stimulation conditions, indicating that these cells (corresponding to DA2) may be more inherently excitable than *Ndnf*+ cells (DA1), which showed the lowest rates of IEG induction. To investigate this possibility, we returned to our snRNAseq data to identify specific pore-forming or auxiliary ion channel genes that might define high- or low-excitability dopamine neurons. A correlation analysis of expression revealed two clusters of ion channel genes that were anti-correlated with one another (**Figure S4K**); when we ran a second correlation on these selected genes along with a composite IEG score for each cell (summed expression of all IEGs, full list), 7 genes had expression positively correlated with IEG activity and 5 genes were anticorrelated with IEG activity (**Figure 4J**).

Genes whose expression was anticorrelated with IEG activation included several potassium channels that are known inhibitors of dopamine neuron activity (*Kcnn3, Kcnd3*, and *Kcnj6*, encoding SK3, Kv4.3, and GIRK2, respectively (Liss et al., 2001, Wolfart et al., 2001, McCall et al., 2017)). The genes positively correlated with IEG expression were also potassium channel subunits, but some of these genes, including *Kcnh7* (Erg3) and *Kcnc2* (Kv3.2) are known to prevent depolarization block or enhance the repolarization needed for fast spiking activity (Rudy and McBain, 2001, Ji et al., 2012), which is consistent with expression in more excitable cells.

In DA neurons, the same ion channel genes that correlated with IEG activity at baseline (in control neurons) were largely correlated with IEG expression following input stimulation (**Figure 4K**). However, expression of these genes was not consistently correlated with IEG activity in GABA, Glu or CoEx cells (**Figure S4L**). Genes anticorrelated with IEG expression showed relatively higher expression in DA1, while genes positively correlated with IEG expression showed higher expression in DA2 (**Figure 4K**). This is consistent with a model of increased baseline excitability and responsiveness in DA2 neurons relative to DA1 neurons, which is reflected in the complement of ion channels expressed in each cell type.

## Discussion

Here we provide a comprehensive transcriptomic dataset of the mouse VTA, characterizing 41,468 total nuclei including 9,336 neurons and 2,006 dopamine neurons. Most previous sequencing studies of this region have excluded non-dopamine cells from analysis, limiting their scope (Poulin et al., 2014, La Manno et al., 2016, Hook et al., 2018, Kramer et al., 2018, Tiklova et al., 2019). A recent snRNAseq study of the rat VTA did include all cell types but described only one dopaminergic population (Phillips et al., 2022). Here we identify 3 molecularly distinct dopamine cell clusters, along with 14 clusters of other cell types, and examine the spatial distribution of 15 marker genes probed in the same VTA tissue. Some of the genes we examined, including *Ndnf, Grp*, and *Pnoc*, had been previously identified in VTA subpopulations (Parker et al., 2019, Poulin et al., 2020), but most are novel markers. It is important to note, however, that few of these genes are perfectly unique markers for a single cell cluster, indicating that future isolation of VTA cell types with fine resolution will almost certainly require combinatorial genetic approaches.

Analysis of the specific pattern of IEGs induced by stimulation of different inputs revealed that cell type, not input, was the strongest predictor of gene activation. A notable exception was that PFC-glutamate stimulation activated a different pattern of IEGs in dopamine neurons than did NAc- or LH-GABA stimulation, even after accounting for overall strength of stimulation.

Intriguingly, this distinction was not observed in GABA, Glu, or CoEx neurons. This indicates a fundamental difference in the transcriptional response in dopamine neurons to direct glutamatergic stimulation versus GABAergic disinhibition and implies that different modes of activation may trigger different long-term changes in dopamine neurons.

Polysynaptic connectivity (i.e. disinhibition or feed-forward excitation) is a major contributor to local VTA circuitry. In addition, many, if not most, glutamatergic and GABAergic inputs co-release neuropeptides that can strongly impact downstream neuronal activity (van den Pol, 2012, Soden et al., 2023). Importantly, the opto-seq approach goes beyond monosynaptic circuit mapping to reveal the net activation of monosynaptic, polysynaptic, and peptidergic signaling. Further investigation using now available genetic tools will allow for dissection of the specific contributions made by fast transmitters and peptides to activation patterns and IEG programs. Highlighting the potential of this approach to uncover novel circuit elements, we found that all three inputs investigated activated at least one GABAergic subcluster, which was unexpected based on the known monosynaptic inhibition of VTA GABA neurons by LH- and NAc-GABA inputs (Nieh et al., 2016, Yang et al., 2018, Soden et al., 2020). The observed activation may be due to direct peptidergic excitation of a subset of GABA neurons, GABA subcluster disinhibition of other GABA subtypes, or potentially an indirect effect of rebound excitation following the termination of stimulation. Distinguishing between these possibilities will require additional dissection of VTA GABA microcircuitry, which will be increasingly possible utilizing the marker genes identified here.

We identified a stimulation-induced pattern of dopamine neuron activation that is consistent with previous analyses of VTA functional heterogeneity. For example, VTA-*Crhr1* neurons, primarily found in our cluster DA1, regulate learning of cue-reward associations, while VTA-*Cck* neurons, found in our clusters DA2 and DA3, regulate motivation (Heymann et al., 2020). Only activation of both populations together drives robust instrumental responding. This is consistent with our finding that only LH-GABA inputs significantly activate DA1 along with DA2/3, as we found previously that these inputs drive the strongest instrumental response (Soden et al., 2020).

Our finding that DA2 neurons showed higher baseline and inducible IEG expression compared to DA1 neurons is also consistent with previous reports of higher firing frequency in medial compared to lateral dopamine neurons (Neuhoff et al., 2002, Lammel et al., 2008). We identified specific ion channel genes that correlated with IEG activity and were differentially expressed between dopamine clusters. Notably nearly all these genes were potassium channel subunits.

The well-studied SK3, Kv4.3, and GIRK2 channels were all associated with high expression in the DA1 cluster and lower IEG activity, consistent with previous reports of higher expression of some of these channels in the lateral VTA (Wolfart et al., 2001, Reyes et al., 2012). Intriguingly, at least two of the channels that correlated with high IEG activity (Erg3 and Kv3.2) have been shown to play a role in enabling fast firing and preventing cells from entering depolarization block (Rudy and McBain, 2001, Ji et al., 2012). Notably, these correlations of expression with IEG activity were unique to dopamine neurons, indicating a cell-type specific role of these ion channels in shaping excitability and responsiveness.

Opto-seq revealed several previously unknown features of VTA input-output connectivity. First, stimulating LH, NAc and PFC inputs led to the recruitment of distinct cellular ensembles that likely reflect downstream behavioral output. Second, different cell types relied on different IEG programs, indicating that measuring expression of a single gene such as *Fos* may be insufficient to fully capture neuronal response dynamics. Third, we can identify genes that correlate with responsivity of specific cell types, allowing us to form testable predictions about gene function and the relationship between cellular excitability and circuit dynamics. Together, these data provide an important advance in our understanding of VTA heterogeneity and demonstrate the utility of combining circuit-specific optogenetics with transcriptomic analysis.

## Acknowledgements

We thank Dasha Krayushkina, Selena Schattauer, and James Allen for assistance with viral production. This work was supported by the University of Washington Royalty Research Fund (M.E.S.) and NIH grants R01 DA054924 (M.E.S), R01 DA044315 and R01 MH104450 (L.S.Z.), R37 DA032750 and R01 DA038168 (G.D.S.), and F31 MH117931 (R.C.S.). This work was also supported by the University of Washington Center of Excellence in Opioid Addiction Research (P30 DA048736) and by the University of Washington W. M. Keck Microscopy Center (S10 OD016240).

## Author contributions

M.E.S. conceptualized the study and designed experiments in consultation with L.S.Z., G.D.S., K.H., and R.C.S. M.E.S. performed experiments with assistance from J.X.Y. and R.C.S. Sequencing data were analyzed by R.C.S., M.E.S., and K.H. *In situ* data were analyzed by M.E.S. and M.L. M.E.S. and R.C.S. prepared figures and wrote the manuscript with input from all authors.

## Methods

### Mice

All procedures were approved and conducted in accordance with the guidelines of the Institutional Animal Care and Use Committee of the University of Washington. Mice were group-housed on a 12-hour light/dark cycle with ad libitum food and water. Approximately equal numbers of male and female mice were used for all experiments. C57BL/6J (strain 000664), *Slc32a1*-Cre (*Vgat*-Cre, strain 028862) and *Slc17a7*-Cre (*Vglut1*-Cre, strain 037512) mice were from Jackson Labs.

### Viruses

AAV1-FLEX-ChR2-YFP and AAV1-FLEX-YFP were produced in-house with titers of 1-3 x10^12^ particles per mL as described (Gore et al., 2013).

### Surgery

Mice were injected at 8-12 weeks of age and recovered for at least 3 weeks prior to experimentation. Mice were anesthetized with isoflurane before and during viral injection and fiber implantation. LH coordinates were M-L: ±1.0, A-P: -1.25, D-V: -5.0. NAc coordinates were M-L: ±0.5, A-P: +1.25, D-V: -4.6. PFC mice received 2 injections per hemisphere at M-L: ±0.35, A-P: +2.1 and +1.4, D-V: -2.0. Values are in mm, relative to bregma. A-P values were adjusted for bregma-lambda distance using a correction factor of 4.21 mm. For Z values the syringe was lowered 0.5 mm past the indicated depth and raised up at the start of the injection. Injection volume was 500 nl. Bilateral optic fibers were implanted over the VTA at M-L: ±0.5, A-P: -3.25, D-V: 4.0.

### Acclimation and optogenetic stimulation

Mice were handled and acclimated to being attached to the patch cord and placed into a clean empty cage for at least 20 min per day for 5 days prior to stimulation. On the stimulation day they received 15 min of 20 Hz, 5 ms blue light stimulation (10 mW).

### Tissue collection and nuclear isolation

Immediately following stimulation mice were anesthetized and perfused with an ice cold solution containing (in mM): 92 NMDG, 2.5 KCl, 1.25 NaH_2_PO_4_, 30 NaHCO_3_, 20 HEPES, 25 glucose, 2 thiourea, 5 Na-ascorbate, 3 Na-pyruvate, 0.5 CaCl_2_, 10 MgSO_4_, pH 7.3-7.4 (Ting et al., 2014). The following inhibitor cocktail was also included (in mM): 5 actinomycin-D, 37.7 anisomycin, 2 kynurenic acid, 0.5 tetrodotoxin. 500 μm coronal VTA sections were cut in the same solution using a VT1000s vibratome (Leica). The VTA from each section was dissected using a fine scalpel blade under a dissecting microscope, transferred to a 1.5 mL tube, snap-frozen on powdered dry ice, and stored at -80°C.

Tissue from 3-4 mice per group was pooled for nuclear isolation, which was conducted as previously described (Hunker and Zweifel, 2020). VTA tissue was dounce homogenized in a buffer containing (in mM): 320 sucrose, 5 CaCl_2_, 3 Mg(C_2_H_3_O_2_)_2_, 10 Tris (pH 7.8), 0.1 EDTA, 0.1 protease inhibitor cocktail, 1 β-mercaptoethanol, 5 actinomycin-D, 37.7 anisomycin, with 0.1% NP40 and 0.2 U/ul RNase inhibitor. Following a 5 min incubation on ice homogenized samples were mixed with an equal volume of 50% iodixanol solution (50% OptiPrep containing (in mM): 5 CaCl_2_, 3 Mg(C_2_H_3_O_2_)_2_, 10 Tris (pH 7.8), 0.1 protease inhibitor cocktail, 1 β-mercaptoethanol, 5 actinomycin-D, 37.3 anisomycin). This mixture was layered on top of an equal volume of 29% iodixanol (OptiPrep) and spun at 7500 RPM for 30 min at 4°C. Pelleted nuclei were resuspended in PBS with 10% BSA and 0.2 U/ul RNAse inhibitor.

### snRNAseq

The concentration of nuclei was determined using a hemocytometer, and was adjusted to 1,000 nuclei/ul. Approximately 10,000 nuclei per experimental group were captured and single-nucleus cDNA libraries were constructed using the Chromium NextGEM platform (10x Genomics) according to manufacturer’s instructions. Libraries were sequenced using an Illumina 4400 HiSeq platform to a target output of ∼ 350,000,000 reads

### Multiplex FISH

For unstimulated experiments, C57BL/6J mice were used. For stimulated experiments, *Vgat*-Cre and *Vglut1*-Cre mice were injected with virus, implanted with fiber-optics, acclimated and stimulated as described above. Immediately following stimulation brains were removed and snap-frozen in crushed dry ice. 20 μm coronal sections of the VTA were made on a CM1950 cryostat (Leica) and mounted on glass slides. Multiple rounds of HCR FISH (Molecular Instruments) were performed on each section, followed by a single round of RNAscope v2 FISH (ACD Bio). Briefly, slides were fixed in 4% PFA followed by ethanol dehydration. Brief protease treatment (3 min, RNAscope Protease III) was used for unstimulated sections only. Each round of HCR was performed according to manufacturer’s instructions, with probes used at a 1:70 dilution. Following amplification, slides were treated with TrueVIEW autofluoresence quenching kit (Vector Laboratories) and mounted with Vectashield Vibrance Antifade mounting medium (Vector Laboratories). Slides were imaged at 20x on a SP8X confocal microscope (Leica). Nuclear DAPI stain was imaged on the first round only. Brightfield images were acquired for all rounds and were used to align images. Following each round of imaging slides were treated with DNase I to remove the probes and allow for another round of probe hybridization, amplification, and imaging. Following the final round of HCR, one round of RNAscope v2 was performed according to manufacturer’s instructions. Brief (3 min) H_2_O_2_ treatment was applied prior to probe incubation.

For unstimulated sections, HCR fluorophores used were 488, 546, and 647, and RNAscope fluorophores (Opal dyes, Akoya Biosciences) used were 520, 570, and 690. For stimulated sections, ChR2-YFP fibers were imaged during the first round. For these experiments HCR fluorophores used were 546, 594, and 647 and RNAscope fluorophores used were 570, 620, and 690.

## Quantification and statistical methods

### General statistics

Statistical tests are reported in each figure legend and were performed using Prism 9 (GraphPad) or R. The Geisser-Greenhouse correction was used to correct for unequal variability of differences in repeated-measures ANOVA tests.

### snRNAseq data analysis

Once sequenced, reads were aligned to a pre-mRNA reference genome using the 10X Genomics’ Cell Ranger v3 pipeline, which provided digital expression matrices that were used for downstream analysis. Clustering, sample integration, cell type identification, and differential gene expression identification were all carried out using the Seurat V3 R package (Stuart et al., 2019).

### Filtering and quality control

General filtering parameters were as previously described (Rossi et al., 2019, Hashikawa et al., 2020). In brief, genes expressed in fewer than 3 cells were removed from the dataset. We next retained cells that possessed between 700-900 and 25,000 UMIs and fewer than 1% mitochondrial read enrichment. Minimum UMI cutoff varied based on sample in attempt to keep final sample quality consistent across groups. Doublets were then computationally predicted and removed by the DoubletDecon R package (DePasquale et al., 2019) using the default parameters.

### Sample integration and clustering

Once putative doublets were omitted, gene transcript counts were scaled for total sequencing depth (factor of 10,000) and then natural-log transformed. We utilized Seurat’s integration algorithm based on canonical correlation analysis (Butler et al., 2018) and mutual nearest neighbor analysis (Haghverdi et al., 2018). Integration anchors were identified for each dataset and transformed by a correction vector to produce a “corrected” expression matrix, on which we performed principal component analysis (PCA). We used the first 30 PCs to identify cell clusters (using Louvain, resolution 0.80) and to plot their proximity in space using Uniform Manifold Approximation and Projection (UMAP). We evaluated the expression of canonical neuronal markers (e.g. *Stmn2* and *Thy1*) to identify cell clusters that were putative neurons, and proceeded to subset these clusters out of the data and performed re-clustering to reveal intra-neuronal heterogeneity. Neurons underwent three clustering iterations in which we removed either low quality cell clusters or potential contaminants. The first iteration was carried out using the same parameters outlined above with the exception of the clustering resolution being limited to 0.40 to prevent over-clustering. Here, we found three cell clusters with low UMI:gene ratio, indicating poor sequencing coverage (Figure S1I-K). We omitted these cell clusters and re-clustered the new neurons again, with the same parameters as the previous iteration. This time, we removed one cluster that had especially poor read coverage in the NAc group, and an additional cluster that appeared to be either microglial contamination or neurons in poor health (enriched expression for *Apoe*). On the final neuron-only iteration, clustering resolution was set to 0.5, yielding a final total of 17 putative neuronal cell types.

### Marker gene identification

To identify potential molecular markers for each cell cluster, we computed the average expression of a given gene in one cluster, and then compared it to the expression of that gene in all other clusters combined. P-values were computed using Wilcoxon rank sum test and corrected for the total number of genes tested (n = 25,501, Bonferroni post-hoc correction).

### IEG analysis

We used a predetermined list of IEGs implicated in the literature, compiled by (Wu et al., 2017). 137 of 139 genes on this list were detectable in our dataset. The short list of 13 commonly studied IEGs was selected prior to any data analysis.

### Correlations

To correlate IEG activation patterns, for each cell class for each stimulus condition we determined the mean expression for each IEG and calculated the log2FC compared to the YFP control group. Genes that were not detected in all conditions were excluded from correlation analysis, which left a total of 96 IEGs from the full list. The cor function in R was used to generate Pearson’s *r* correlation coefficient values and the heatmap.2 function was used to plot the correlation.

To correlate ion channel expression we identified ion channel genes that were expressed in at least 25% of cells in at least one of the three DA clusters. We determined the expression of each of these genes in each cell in the DA clusters across all 4 groups and the cor and heatmap.2 functions in R were used to generate and plot the correlation of gene expression. From this analysis we selected one cluster of 5 genes and one cluster of 7 genes that appeared most anticorrelated with one another. We repeated the correlation analysis on this subset of genes, also including an IEGscore calculated for each cell as the sum of the expression of all IEGs (full list).

### HCR data analysis

Images were aligned using HiPlex Image Registration software (ACDBio) with brightfield images as a reference. In some cases, when automated alignment failed, images were manually aligned using Photoshop (Adobe). For some probes, non-specific punctate background signal was removed using the Remove Outliers function in ImageJ. Image analysis was performed using Halo software v3.2 (Indica Labs). Copy number thresholds and parameters for detecting puncta and intensity of each gene were manually adjusted for each section for each animal. Cell type abundance, spatial distributions, and correlations of IEG activation with ChR2 fibers were calculated using custom R scripts. For ChR2-YFP fiber analysis each section was divided into a grid of approximately 150x150 μm squares and all cells expressing *Th, Gad2*, or *Vglut2* were included in analysis. ChR2-YFP fiber intensity was calculated as the average YFP fluorescence per cell in each square, as detected by Halo.

## Data and code availability

Data and code used in this paper will be made publicly available upon publication of the manuscript in a peer-reviewed journal.

