## Supplementary Figures for "Opto-seq reveals input-specific immediate early gene induction in ventral tegmental area cell types"

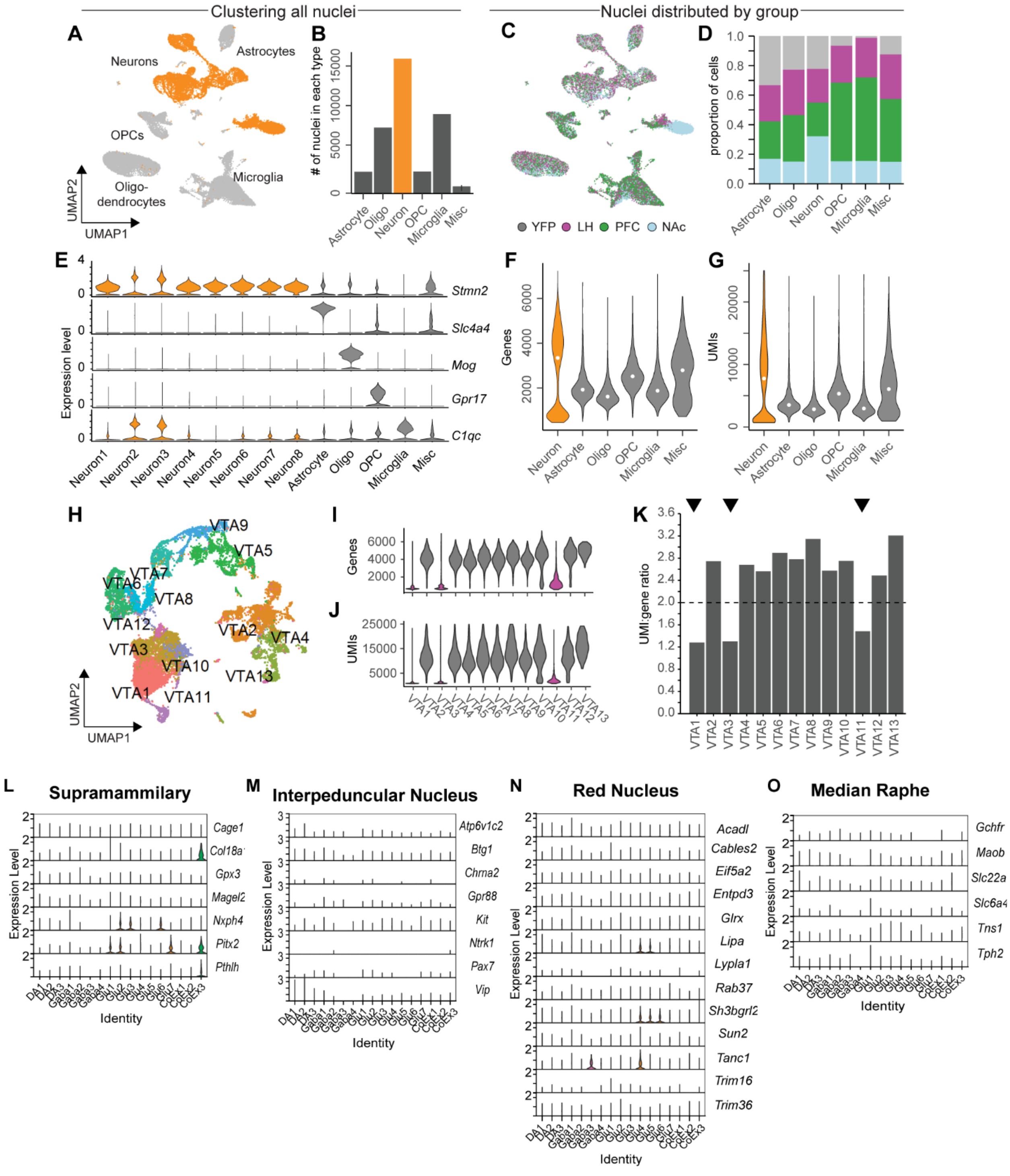


***Figure S1. A)** UMAP plot of all nuclei, identified by expression of cell-type markers. **B)** Number of nuclei of each type. **C)** Distribution of cells from different stimulus conditions across clusters. **D)** Proportion of cells of each type from each stimulus condition. **E)** Violin plot showing expression of cell-type markers. Note that neuron clusters 2 and 3 showed substantial expression of microglial markers along with neuronal markers; these cells were included in the initial reclustering of neurons but were largely eliminated during further quality control steps (see Methods). **F-G)** Violin plots depicting number of genes*

*and unique molecular identifiers (UMIs) per nuclei. Note the bimodal distribution in neurons: nuclei with low gene and UMI counts were largely eliminated during further quality control steps (see Methods and panels I-K).* ***H)*** *UMAP plot showing initial reclustering of neurons.* ***I-J)*** *Violin plots depicting genes and UMIs per nuclei in neuronal subclusters.* ***K)*** *UMI/gene ratio for neuronal subclusters. Subclusters marked with triangles were deemed to be low quality nuclei and were not included in further analysis.* ***L-O)*** *Violin plots depicting expression of genes identified through the Allen Brain Atlas as being enriched in the indicated brain regions relative to the VTA.*

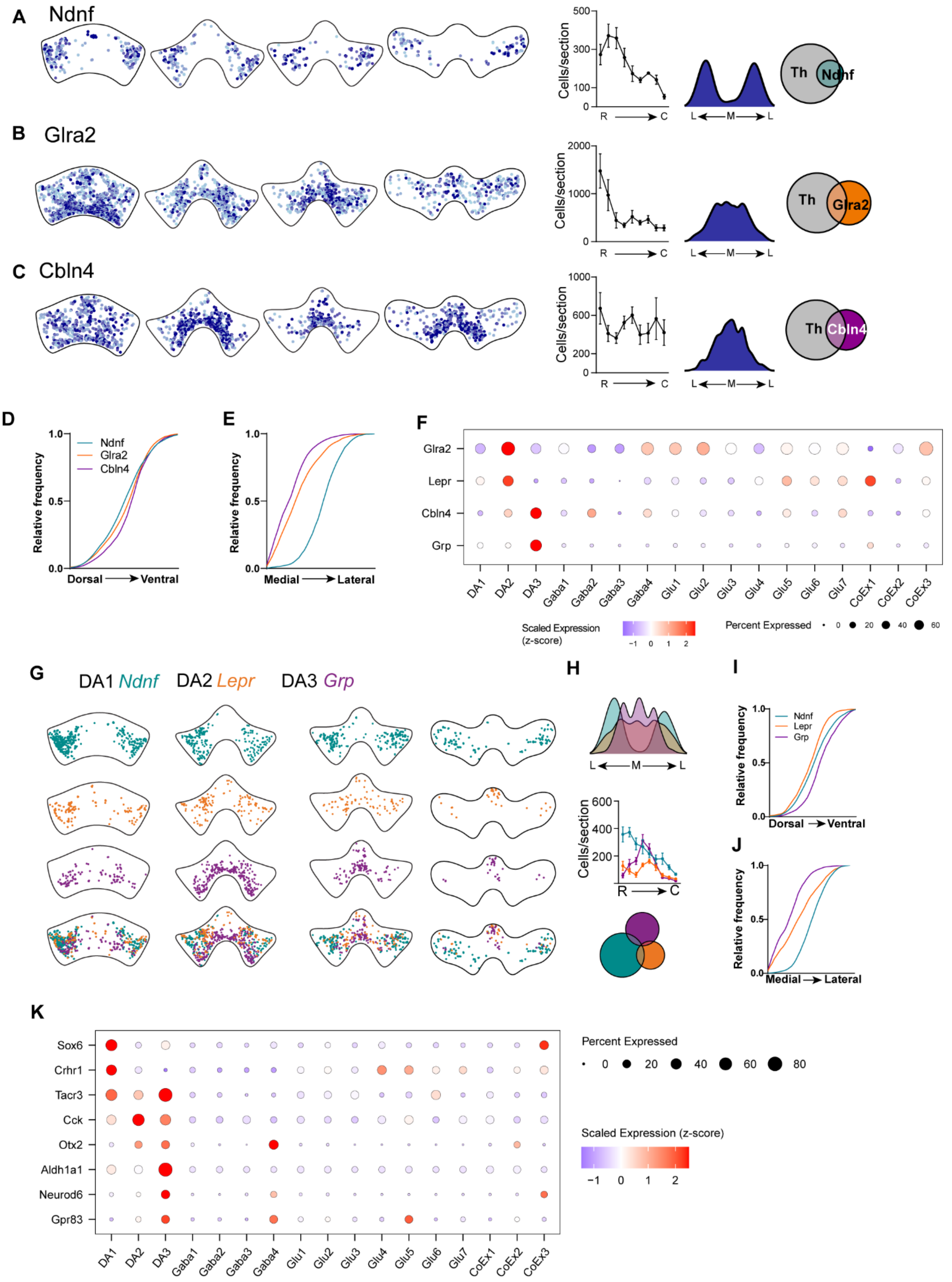

A Ndnf
B Glra2
C Cbln4
Cells/section
R → C
L ← M → L
Th
Ndnf
Glra2
Cbln4
D
E
F
Relative frequency
Dorsal → Ventral
Medial → Lateral
Glra2
Lepr
Cbln4
Grp
DA1 DA2 DA3 Gaba1 Gaba2 Gaba3 Gaba4 Glu1 Glu2 Glu3 Glu4 Glu5 Glu6 Glu7 CoEx1 CoEx2 CoEx3
Scaled Expression (z-score)
Percent Expressed
G
DA1 Ndnf DA2 Lepr DA3 Grp
H
I
J
Grp
K
Sox6
Crhr1
Tacr3
Cck
Otx2
Aldh1a1
Neurod6
Gpr83

***Figure S2. A, B, and C)*** *From left to right: Representative expression plots of all Ndnf+, Glra2+, or Cbln4+ cells (not just those that are Th+); Average number of cells per section, rostral (R) to caudal (C), n=4 mice, 9 sections, mean±SEM; Probability density plot showing medial (M) to lateral (L) distribution of cells, data pooled from 4 mice; Euler diagram showing overlap with Th, data pooled from 4 mice.* ***D)*** *Cumulative distribution of Th+ cells co-labeled with indicated subcluster markers from dorsal to ventral across 9 sections from each of 4 mice. Mann-Whitney test for all comparisons (Ndnf vs. Glra2, Ndnf vs. Cbln4, Glra2 vs. Cbln4): P<0.0001.* ***E)*** *Cumulative distribution of Th+ cells co-labeled with indicated subcluster markers from medial to lateral across 9 sections from each of 4 mice. Mann-Whitney test for all comparisons (Ndnf vs. Glra2, Ndnf vs. Cbln4, Glra2 vs. Cbln4): P<0.0001.* ***F)*** *Dot plot showing relative expression (z score) of original DA2 and DA3 markers Glra2 and Cbln4, and alternative markers Lepr and Grp.* ***G)*** *Representative expression plots of Th+ cells that were also positive for the indicated markers.* ***H)*** *Top: Probability density plots showing medial (M) to lateral (L) distribution of cells. Data pooled from 4 mice. Middle: Average number of cells per section, rostral (R) to caudal (C) (N=4 mice, 9 sections, mean±SEM). Bottom: Euler diagram showing overlap between Ndnf, Lepr, and Grp. Data pooled from 4 mice.* ***I-J)*** *Cumulative distributions of Th+ cells co-labeled with indicated subcluster markers from dorsal to ventral (**I**) or medial to lateral (**J**). Data pooled from 4 mice. Mann-Whitney test for all comparisons (Ndnf vs. Lepr, Ndnf vs. Grp, Lepr vs. Grp): P<0.0001.* ***K)*** *Dot plot showing relative expression (z-score) of select dopamine neuron subtype markers identified by previous studies.*

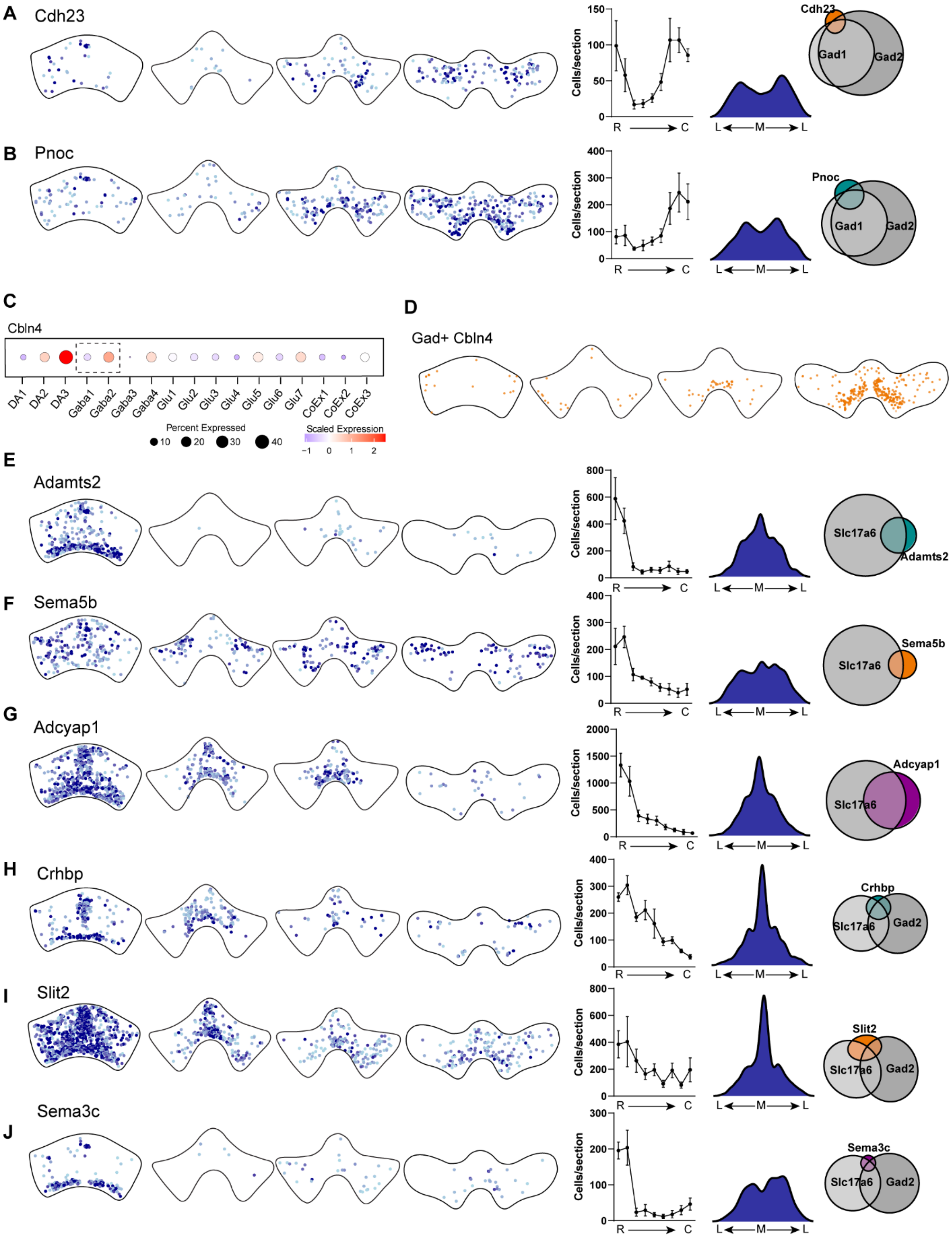

A
Cdh23
Cells/section
150
100
50
0
R
C
L
M
L
Cdh23
Gad1
Gad2
B
Pnoc
400
300
200
100
0
Pnoc
Gad1
Gad2
C
Cbln4
DA1
DA2
DA3
Gaba1
Gaba2
Gaba3
Gaba4
Glu1
Glu2
Glu3
Glu4
Glu5
Glu6
Glu7
CoEx1
CoEx2
CoEx3
Percent Expressed
10
20
30
40
Scaled Expression
−1
0
1
2
D
Gad+ Cbln4
E
Adamts2
800
600
400
200
0
Slc17a6
Adamts2
F
Sema5b
400
300
200
100
0
Sema5b
Slc17a6
G
Adcyap1
2000
1500
1000
500
0
Adcyap1
Slc17a6
H
Crhbp
400
300
200
100
0
Crhbp
Slc17a6
Gad2
I
Slit2
800
600
400
200
0
Slit2
Slc17a6
Gad2
Sema3c
J
300
200
100
0
Sema3c
Slc17a6
Gad2

***Figure S3. A-B)*** *From left to right: Representative expression plots of all Cdh23+ or Pnoc+ cells; Average number of cells per section, rostral (R) to caudal (C), n=4 mice, 9 sections, mean±SEM; Probability density plot showing medial (M) to lateral (L) distribution of cells, data pooled from 4 mice; Euler diagram showing overlap with indicated cell-type markers, data pooled from 4 mice.* ***C)*** *Dot plot showing scaled expression of Cbln4 across subclusters, highlighting expression in GABA2 cluster* ***D)*** *Representative expression plots of Cbln4+ cells that are also Gad1 or Gad2 positive.* ***E-J)*** *Plots as in (**A**), for indicated genes.*

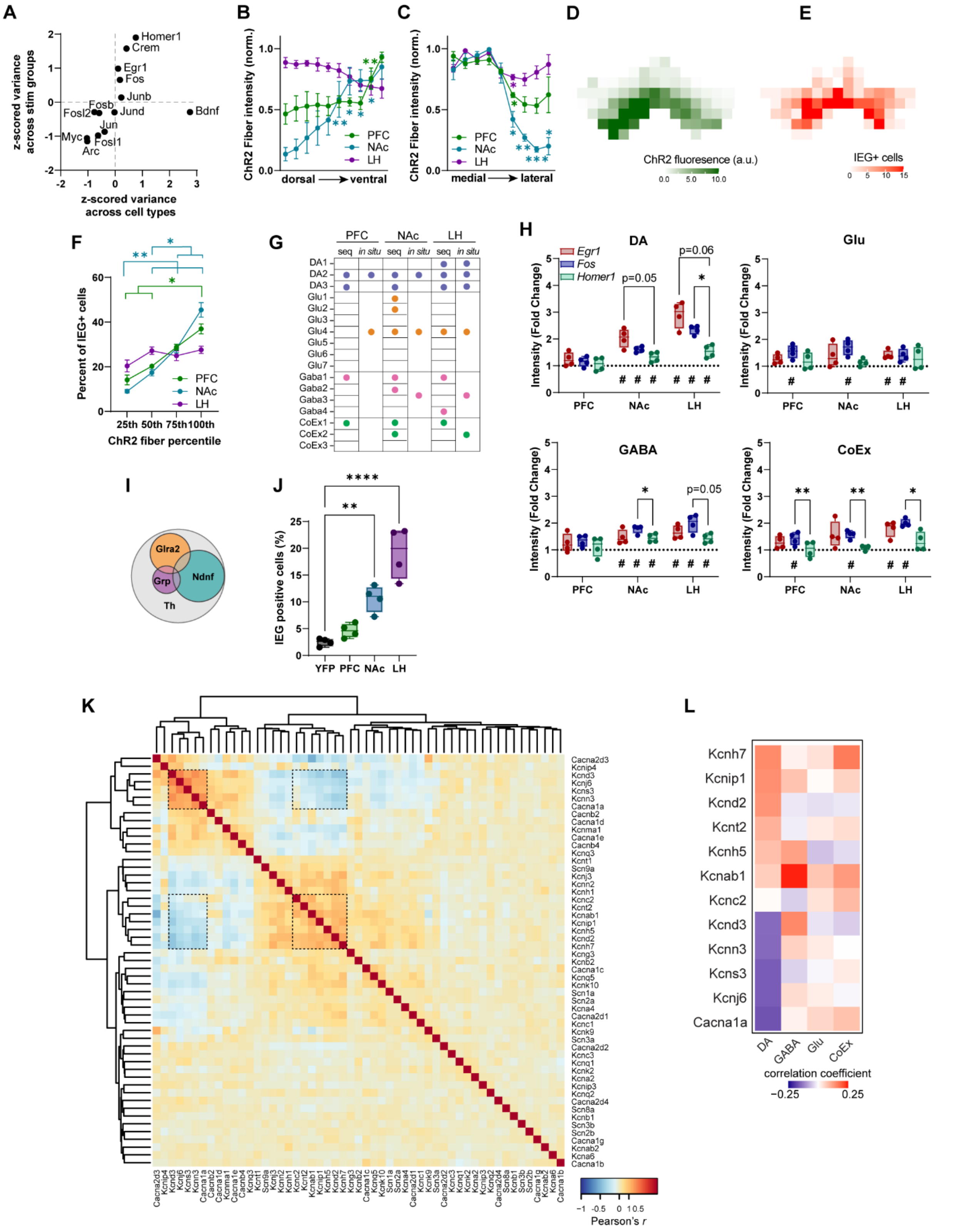
A
z-scored variance across stim groups
z-scored variance across cell types
Homer1
Crem
Egr1
Fos
Junb
Fosb
Fosl2
Jund
Bdnf
Jun
Myc
Fosl1
Arc
B
ChR2 Fiber intensity (norm.)
dorsal ventral
PFC
NAc
LH
C
medial lateral
D
ChR2 fluoresence (a.u.)
E
IEG+ cells
F
Percent of IEG+ cells
ChR2 fiber percentile
25th 50th 75th 100th
G
PFC NAc LH
seq in situ
DA1 DA2 DA3
Glu1 Glu2 Glu3 Glu4 Glu5 Glu6 Glu7
Gaba1 Gaba2 Gaba3 Gaba4
CoEx1 CoEx2 CoEx3
H
DA
Glu
GABA
CoEx
Egr1
Fos
Homer1
Intensity (Fold Change)
p=0.05
p=0.06
I
Glra2
Grp
Ndnf
Th
J
IEG positive cells (%)
YFP PFC NAc LH
K
Cacna2d3 Kcnip4 Kcnd3 Kcnj6 Kcns3 Kcnn3 Cacna1a Cacnb2 Cacna1d Kcnma1 Cacna1e Cacnb4 Kcnq3 Kcnt1 Scn9a Kcnj3 Kcnn2 Kcnh1 Kcnc2 Kcnt2 Kcnab1 Kcnip1 Kcnh5 Kcnd2 Kcnh7 Kcng3 Kcnb2 Cacna1c Kcnq5 Kcnk10 Scn1a Scn2a Kcna4 Cacna2d1 Kcnc1 Kcnk9 Scn3a Cacna2d2 Kcnc3 Kcnq1 Kcnk2 Kcna2 Kcnip3 Kcnq2 Cacna2d4 Scn8a Kcnb1 Scn3b Scn2b Cacna1g Kcnab2 Kcna6 Cacna1b
Pearson's r
L
Kcnh7 Kcnip1 Kcnd2 Kcnt2 Kcnh5 Kcnab1 Kcnc2 Kcnd3 Kcnn3 Kcns3 Kcnj6 Cacna1a
DA GABA Glu CoEx
correction coefficient
−0.25 0.25

***Figure S4. A)*** *Z-scored variance of percent of cells expressing each IEG between cell-types (x) and between stimulus conditions (y).* ***B)*** *Mean ChR2 fiber intensity of all cells from all sections, binned across the dorso-ventral axis, normalized to the maximum bin for each mouse (N=4 mice/group, 2-way RM ANOVA $F_{(18, 81)}$=12.13, P<0.0001, Tukey's multiple comparisons within group, *P<0.05, **P<0.01 labeled for any point in ventral half significantly different from any point in dorsal half).* ***C)*** *Mean ChR2 fiber intensity of all cells from all sections, binned across the medial-lateral axis, normalized to the maximum bin for each mouse (N=4 mice/group, 2-way RM ANOVA $F_{(16, 72)}$=7.836, P<0.0001, Tukey's multiple comparisons within group, *P<0.05, **P<0.01, ***P<0.001 labeled for any point in lateral half significantly different from any point in medial half).* ***D-E)*** *Example from a NAc mouse of ~150x150 um grid applied to 1 VTA section and fiber intensity (D) and number of IEG+ cells (E) calculated for each division.* ***F)*** *Percent of all IEG+ cells found in grid divisions separated into quartiles by ChR2 fiber fluorescence intensity (N=4 mice/group, mean±SEM. 2-way RM ANOVA, interaction $F_{(6, 27)}$=9.249, P<0.0001; Tukey's multiple comparisons *P<0.05, **P<0.01).* ***G)*** *Summary of clusters significantly activated (P<0.05) compared to YFP control in sequencing and in situ studies. For in situ data Ndnf, Glra2, and Grp were used to isolate DA clusters, but subcluster markers were not tested for Glu, GABA, or CoEx neurons so these groups are merged.* ***H)*** *Fold-change in fluorescence intensity of cells positive for each IEG (compared to mean of YFP group) in each cell type. (N=4 mice/group. One-sample t-test for each gene to determine if significantly different from 1 (control): #P<0.05. Two-way RM ANOVA: DA interaction: $F_{(4,18)}$=4.515, P=0.0106; GABA effect of input: $F_{(2,9)}$=8.347, P=0.0089, effect of gene: $F_{(1.328, 11.95)}$=6.219, P=0.0218; CoEx effect of input: $F_{(2,9)}$ = 9.878, P=0.0054, effect of gene: $F_{(1.134,10.21)}$=10.21, P=0.0080. Tukey's multiple comparisons, *P<0.05, **P<0.01).* ***I)*** *Euler diagram showing proportion of Th+ cells expressing indicated markers. Data pooled from 16 mice.* ***J)*** *Percent of cells expressing Th but no subgroup marker that are positive for one or more IEG (N=4 mice/group, 1-way ANOVA, $F_{(3, 12)}$=28.34, P<0.0001, Dunnett's multiple comparisons, each stim group versus control: **P<0.01, ****P<0.0001).* ***K)*** *Heatmap showing the correlation coefficient (r) of ion channel gene expression in dopamine neuron clusters. Highlighted regions show genes featured in Figure 4J.* ***L)*** *Heatmap of correlation coefficient of each ion channel gene with the IEG score, separated by cell type.*
